# A parasitoid serpin gene that disrupts host immunity shows adaptive evolution of alternative splicing

**DOI:** 10.1101/2023.03.28.534536

**Authors:** Zhichao Yan, Qi Fang, Lei Yang, Shan Xiao, Jiale Wang, Gongyin Ye

## Abstract

Alternative splicing (AS) is a major source of protein diversity in eukaryotes, but less is known about its evolution compared to gene duplication (GD). How AS and GD interact is also largely understudied. By constructing the evolutionary trajectory of a serpin gene PpSerpin-1 (*Pteromalus puparum* serpin 1) in parasitoids and other insects, we found that both AS and GD jointly contribute to serpin protein diversity. These two processes are negatively correlated and show divergent features in both protein and regulatory sequences. Furthermore, parasitoid wasps exhibit higher numbers of serpin protein/domains than nonparasitoids, resulting from more GD but less AS in parasitoids. Nevertheless, PpSerpin-1 shows an exon expansion of AS compared to other parasitoids. We find that several isoforms of PpSerpin-1 are involved in the wasp immune response, have been recruited to both wasp venom and larval saliva, and suppress host immunity. In summary, we report the differential features of AS and GD in the evolution of insect serpins and their associations with the parasitic life strategy, and we provide an example of how a parasitoid serpin gene adapts to parasitism through AS.

## Introduction

Alternative splicing (AS) is an ubquitious regulatory process of transcription in animals, plants and fungi^1,2^. Through differential inclusion/exculsion of exons and introns, AS produces multiple variant proteins from a single gene^1,2^. AS and gene duplication (GD) are important sources of genetic innovation and protein diversity^3–6^. Compared to GD, relatively little is known about the evolution of and role of AS in adaptation^3,7,8^. Moreover, the relationship and difference between these two evolutionary processes, AS and GD, are largely understudied^4,9,10^.

Pathogens and parasites recruit diverse virulence genes to overcome their hosts and adapt to coevolution with hosts^11–14^. For example, to ensure successful parasitism, parasitoid wasps often inject venom to manipulate their hosts’ immunity, metabolism, development and even behavior^14,15^. Driven by frequent host shifts and the arms race between parasitoid wasps and their hosts, parasitoid venoms show rapid compositional turnover and high sequence evolutionary rates^14,16–18^. Evolutionary processes for parasitoid venom gene recruitment include GD^19,20^ ^21^, AS^22,23^, lateral gene transfer^24,25^, and single-copy gene co-option for venom functions^17,26^.

Serpins (<u>ser</u>ine <u>p</u>rotease <u>in</u>hibitors) are a widely distributed protein superfamily in all kingdoms of life^27–29^. Serpins contain similar structures with three β-sheets, 7–9 α-helices and an exposed reactive center loop (RCL) on the C-terminus, which determines serpin activity and specificity. Most serpins are inhibitors that irreversibly inhibit target enzymes by conformational change^30^, where the hinge region of RCL acts as a “spring” and inserts into β-sheets^31,32^. Through their inhibitory activities, serpins play central roles in proteolytic cascades, e.g., coagulation and inflammation in mammals and the prophenoloxidase (PPO) casecade and Toll pathway in insects^28,29^.

In parasitoid wasps, serpins are a common venom component and have been reported in numerous parasitoid venoms^17, 20–23, 33–41^. Both AS and GD have been reported in the recruitment of serpin into parasitoid venoms^21,22^. Previously, we isolated a venom serpin protein from *Pteromalus puparum*, a generalist parasitoid wasp that parasitizes pupae of several butterfly species^22^. This venom serpin is produced by the PpSerpin-1 gene through AS and suppresses the host’s melanization immunity^22^. In contrast, extensive GD of serpin was reported in the venom apparatus of a parasitoid wasp, *Microplitis mediator*^21^. Therefore, parasitoid venom serpins may provide a promising model for comparative studies on the evolution and ecological adaptation of AS and GD.

Here, we analyzed the evolutionary trajectory of PpSerpin-1 and compared the different features of AS and GD in serpin evolution and their associations with the parasitic life strategy. In addition, we demonstrate that PpSerpin-1 is involved in wasp’s immune responses and has been recruited to both wasp venom and larval saliva, to regulate host immunity.

## Results

### AS contributes to serpin protein diversity with GD and domain duplication in insects

Utilizing the accumulated transcriptomic data, we found that the PpSerpin-1 gene has alternate splicing, with two different N-terminal and 21 C-terminal AS forms (Fig. 1a, S1). At the N-terminus, one form was predicted to be secreted extracellularly with a signal peptide, while the other was predicted to localize intracellularly without a signal peptide (Fig. 1b). At the C-terminus, AS occurs at the end of the serpin hinge region with an extra nucleotide G (Fig. 1b, S1).

**Figure 1.**
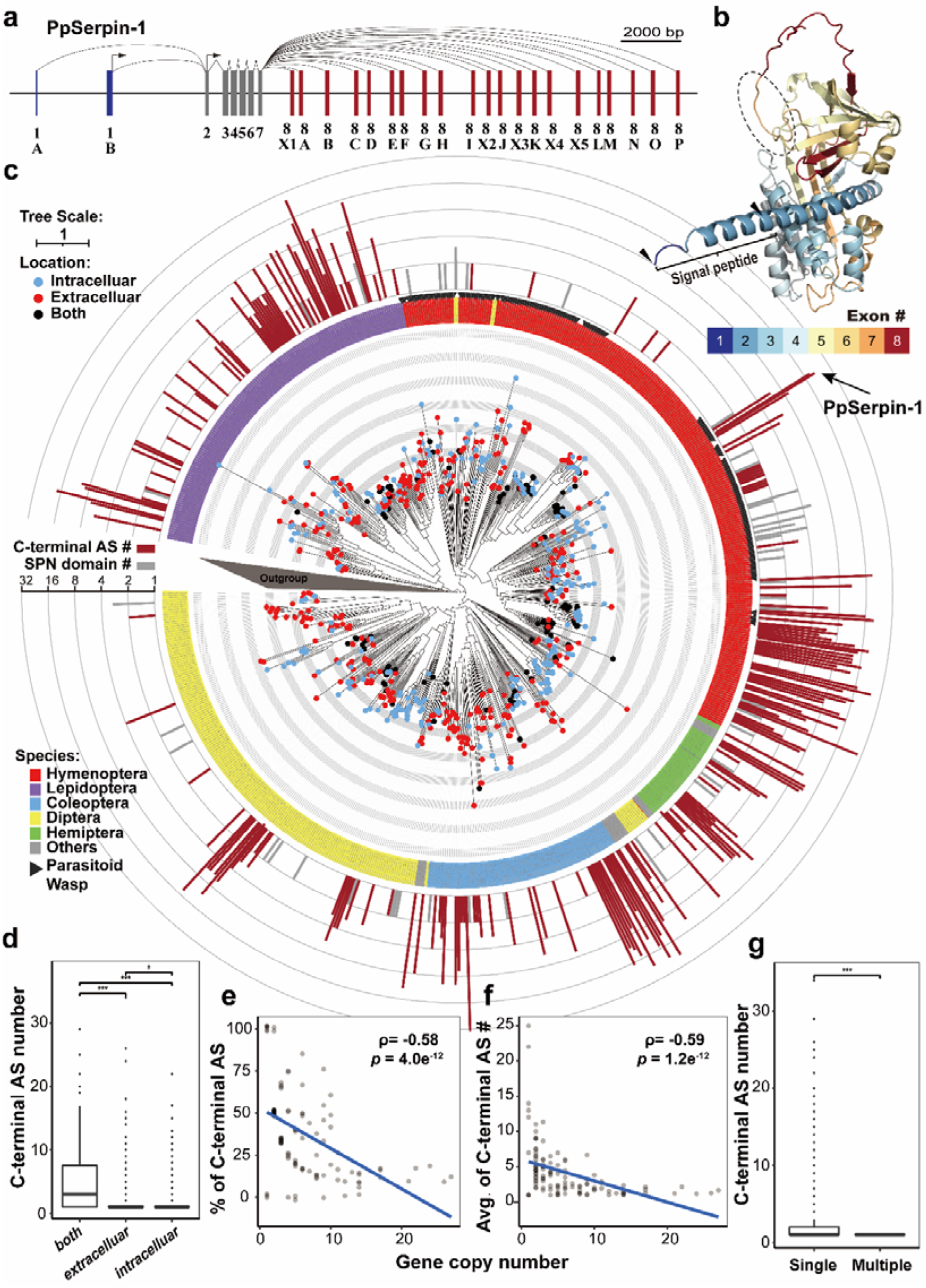
Alternative splicing (AS), gene duplication (GD) and domain duplication contribute to serpin protein diversity in insects. (a) Gene structure of PpSerpin-1. Gray indicates constitutive exon. Blue and red indicate N-terminal and C-terminal alternative exons, respectively. (b) Protein structure of PpSerpin-1F predicted by AlphaFold2. PpSerpin-1F is the longest protein isoform of the PpSerpin-1 gene and was chosen as the representative. The gradient colors from blue to red indicate different exons. The bracket indicates the signal peptide. Black arrows indicate alternative translation start sites. The dashed circle indicates the hinge region of PpSerpin-1F. (c) Phylogenetic tree of serpin in insects. The different colors on the labels indicate different insect orders. Black arrows indicate parasitoid wasps. The red and blue dots on branch terminals indicate the presence (extracellular) and absence (intracellular) of the signal peptide, respectively. Black dots on branch terminals indicate N-terminal AS with both extracellular and intracellular isoforms. The red bars indicate the log2 transformed numbers of C-terminal AS events. The gray bars indicate the log2 transformed numbers of serpin domains within a gene. (d) Relationship between the number of C-terminal AS events and gene localization. (e) Relationship between the percentage of C-terminal alternative spliced genes and serpin gene copy number. (f) Relationship between the number of C-terminal AS events and serpin gene copy number. (g) Relationship between the number of C-terminal AS events and sperin domain number. *** *p* < 0.001; * *p* < 0.05.

To construct the phylogeny of the PpSerpin-1 gene, a homology search was performed against the insect Refseq_protein database in NCBI using the constitutive region of PpSerpin-1 (for details, see Materials and Methods), resulting in a total of 1,230 matched genes. Phylogenetic analyses showed that the C-terminal AS of serpin genes was clustered into one clade with 731 genes but was relatively rare in the outgroup (Fig. 1c). The following analyses were all focused on this clade with enriched C-terminal AS.

Both N-terminal and C-terminal AS are widespread in insects (Fig. 1c). Out of 731 genes in this clade, 261 genes (35.7%, not including PpSerpin-1) have either N-terminal or C-terminal AS. At the N-terminus, 153 out of 731 (21.0%) had N-terminal AS, with 115 out of 153 having both extracellular and intracellular forms. These genes with both extracellular and intracellular forms at the N-terminus had more C-terminal AS than genes with only one N-terminal form, either extracellular or intracellular (Fig. 1d; MannCWhitney U test (MWUT); both comparisons: *p* < 0.001). At the C-terminus, 192 out of 731 (26.3%) genes had C-terminal AS producing 1406 serpin proteins, with an average of 7.32 serpin proetins per gene. Notably, the number of C-terminal AS events showed rapid fluctuation, suggesting a high turnover rate of exon gain and loss (Fig. 1c).

Together, AS, GD and domain duplication within genes contribute to serpin protein diversity. Forty-five genes have multiple serpin domains, producing 105 serpin domains with an average of 2.33 serpin domains per gene. The remaining 494 genes, which have neither C-terminal AS nor multiple serpin domains, likely originated by GD. Taken together, AS, GD and domain duplication account for 70.1%, 24.6% and 5.2% of the total number of 2005 serpin proteins/domains, respectively. Moreover, AS is negatively correlated with GD and domain duplication. The number of duplicated serpin gene copies in a species is inversely correlated with the percentage of C-terminal AS (Fig. 1e; Spearman correlation; ρ = −0.58, *p* = 4.0e^-12^) and with the means of C-terminal variants (Fig. 1f; Spearman correlation; ρ = −0.59, *p* = 1.2e^-12^). Both correlations held after phlogentic correction (both tests: *p* < 0.001). In addition, AS and domain duplication within genes are mutually exclusive, as no multiple-serpin gene contains C-terminal AS (Fig. 1g; MWUT; single- vs multiple-domain serpin: *W* = 19755, *p* = 4.8e^-5^).

### AS genes show sequence features divergent from non-AS genes

Both AS and GD are major sources of protein diversity. To compare their differences in gene sequence characteristics, we divided the single-domain serpins into genes with C-terminal AS (referred to as the AS gene set) and those without (referred to as the non-AS gene set). Here, we focused on C-terminal but not N-terminal AS, as N-terminal AS often changes protein localization but not the mature serpin protein sequence.

For the AS gene set, the splicing positions of C-terminal AS occurred mainly near the end of the hinge region (GSEAAAVT in PpSerpin-1) (Fig. 2a). Considering the last nucleotide at the end of the hinge region as position 0, 161 out of 192 (83.9%) AS genes splice at position +1 and 19 (9.9%) at position +4. The AS gene set is more likely to splice at position +1 than the non-AS gene set (Fig. 2a; χ*²* = 35.81, *p* = 0.00001). Both AS and non-AS gene sets have G|GTAAGT and TTNCAG|N sequence motifs near C-terminal splicing sites (Fig. 2b). Position 0 (at codon position 3), +5 (at codon position 2) and +11 (at codon position 2) are more inclined to be G, T and T in the AS gene set than in the non-AS gene set, respectively (Fig. 2b; *p* < 0.05).

**Figure 2.**
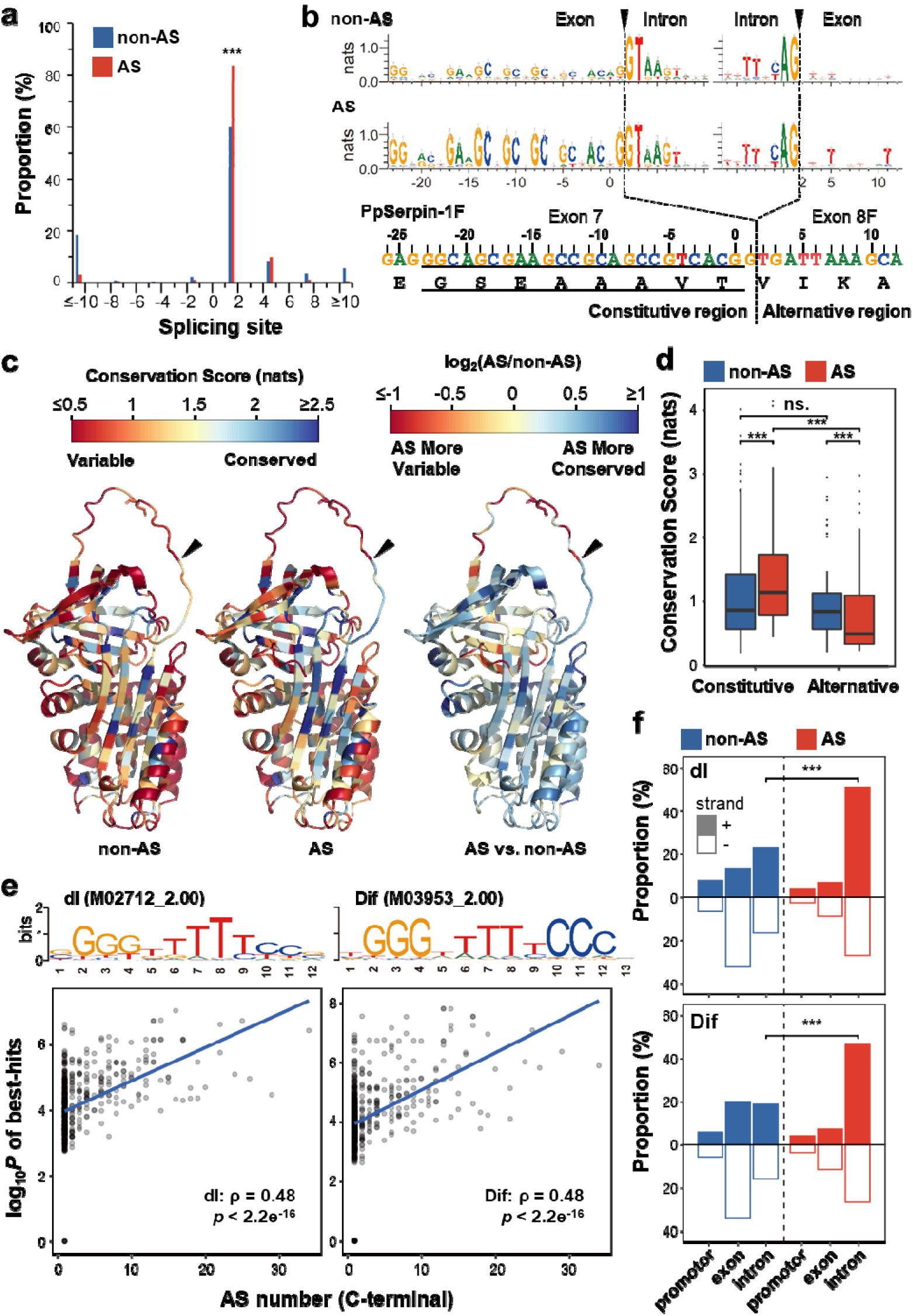
AS genes show divergent sequence features from non-AS genes. (a) Distribution of splicing sites on the AS and non-AS gene sets. The AS gene set represents single-domain serpin genes with C-terminal AS, and the non-AS gene set represents single-domain serpin genes without C-terminal AS. The end of the hinge region is noted as position 0. **(b)** Sequence logos of AS and non-AS gene sets. The dashed lines indicate splicing positions. The underline indicates the hinge region. **(c)** Predicted protein structure of PpSerpin-1F with mapped conservation scores of AS and non-AS gene sets. **(d)** Comparisons of conservation scores between AS and non-AS genes. **(e)** Correlation of C-terminal AS number and probability of dl and Dif binding motif hits. **(f)** Binding motif hit distribution of dl and Dif on AS and non-AS genes. + and - indicate coding and noncoding strands, respectively. *** *p* < 0.001; ns. : *p* > 0.05.

For the protein sequence, we divided the serpin proteins into constitutive (present in all isoforms) and C-terminal alternative spliced regions based on the end of the hinge region (Fig. 2b). Conservation scores were estimated based on alignments and then mapped to PpSerpin-1F as the reference (Fig. 2c). In the constitutive region, the AS gene set is more conserved than the non-AS gene set (Fig. 2c, 2d; Wilcoxon signed-rank test (WSRT); *V* = 985, *p* < 2.2e^-16^). Conversely, in the alternative region, the AS gene set is more variable than the non-AS gene set (Fig. 2c, 2d; WSRT: V = 1125, *p* = 1.5e^-05^). To exclude the effect of reference sequence selection, we also used non-AS genes within the clade (Dmel_SPN55B, NP_524953.1) or in the outgroup (Dmel_SPN27A, NP_001260143.1) as reference sequences, and the conclusions held regardless of reference selection (all comparisons: *p* <0.001).

Furthermore, we compared the regulatory sequences of the AS and non-AS gene sets. For RNA-binding motifs, no motifs were found enriched in the AS gene set compared to the non-AS gene set (*p* > 0.05), which may be due to the general short length of RNA-binding motifs and lack of statistical potency, as previously reported^42^. For binding motifs of transcription factors, two motifs are enriched in the AS gene set compared to non-AS. They are M02712_2.00 dl (dorsal) (Fig. 2e; *p* = 0.010, enrichment ratio 8.01) and M03953_2.00 Dif (Dorsal-related immunity factor) (*p* = 0.015, enrichment ratio 7.39). dl and Dif are both transcription factors involved in the Toll pathway^43^. Compared to shuffled random sequences, dl and Dif binding motifs are significantly enriched in the AS gene set (dl: *p* = 3.33e^-7^; Dif: *p* = 7.70e^-6^) but not the non-AS gene set (*p* > 0.05). In addition, the probabilities of the presence of dl and Dif binding motifs are significantly correlated with the numbers of C-terminal AS (Fig. 2e; Spearman correlation; dl: ρ = 0.48, *p* < 2.2e^-16^, dif: ρ = 0.48, *p* < 2.2e^-16^). These correlations are significantly higher than correlations using shuffled sequences, which have the same lengths as the true gene sequences (both comparisions: *p* < 0.0001). These results suggest that genes with more C-terminal AS are more likely to contain dl and Dif binding motifs, and this is not simply due to their longer sequence lengths.

We further asked how these dl and Dif binding motifs are distributed in sequences. We defined 6 region categories according to their strands (on coding or noncoding strands) and location (in promoter, exon or intron regions). Most of the best-hits are in the intron region of the coding strand for the AS gene set, while most of the best-hits are in the exon region of the noncoding strand for the non-AS gene set (Fig. 2f). Best-hits of dl and Dif binding motifs are more likely to locate in the intron region of coding strand of the AS gene set than that of the non-AS gene set (Fig. 2f; dl: χ*²* = 50.40, *p* < 0.00001; dif: χ*²* = 55.38, *p* < 0.00001) and shuffled control sequences of the AS gene set (dl: χ*²* = 10.8984, *p* = 0.000962; dif: χ*²* = 7.9206, *p* = 0.004888). In addition, when we restricted the motif search to one of the six region categories, the correlations between the probabilities of the presence of dl and Dif binding motifs and the numbers of C-terminal AS were significantly higher than correlations using shuffled sequences in the intron region of the coding strand but not in the other five region categories (dl: *z* = 2.7, *p* = 0.0065; dif: *z* = 4.4, *p* < 0.0001; all other comparisons: *p* > 0.05). Therefore, we conclude that dl and Dif binding motifs tend to be enriched in the intron region of the coding strand within the AS gene set.

### Parasitoid wasps employ fewer AS in serpin, but PpSerpin-1 is an expansion of the AS exon number

We further investigated the potential relationship between AS and the parasitic life strategy. Higher numbers of total serpin protein/domain per species were found in parasitoid wasps than in nonparasitoid hymenopterans (Fig. 3a; *W* = 370, *p* = 0.0026). Compared to nonparasitoid hymenopterans, more GD (Fig. 3b; MWUT; *W* = 434, *p* = 5.5e^-6^) and domain duplication (Fig. S2; MWUT; average: *W* = 362.5, *p* = 2.1e^-4^; proportion: W = 350, p = 8.1e-4) but less AS (Fig. 3c, 3d; MWUT; average: *W* = 96, *p* = 0.0022; proportion: *W* = 96, *p* = 0.0017) is utilized in parasitoid wasps to produce serpin protein/domain diversity. Consistent with this, gene expansions in parasitoid wasps are generally more recent (Fig. S3a; MWUT; *W* = 11429, *p* = 3.0e^-6^) with a higher proportion of genus-specific and family-specific expansions (Fig. 3e; genus-specific: χ*²* = 14.09, *p* = 1.74e^-4^; family-specific: χ*²* = 10.49, *p* = 1.20e^-3^) than those in nonparasitoid hymenopterans. Conversely, exon expansions in parasitoids are generally more ancient (Fig. S3b; MWUT; *W* = 16338, *p* = 1.25e^-4^) with a lower proportion of lineage-specific exon expansions (Fig. 3f; χ*²* = 20.14, *p* < 0.00001) than those in nonparasitoid hymenopterans. Patterns are similar when comparing parasitoid wasps with all nonparasitoid insects (not limited to Hymenoptera) (all comparisons: *p* < 0.05).

**Figure 3.**
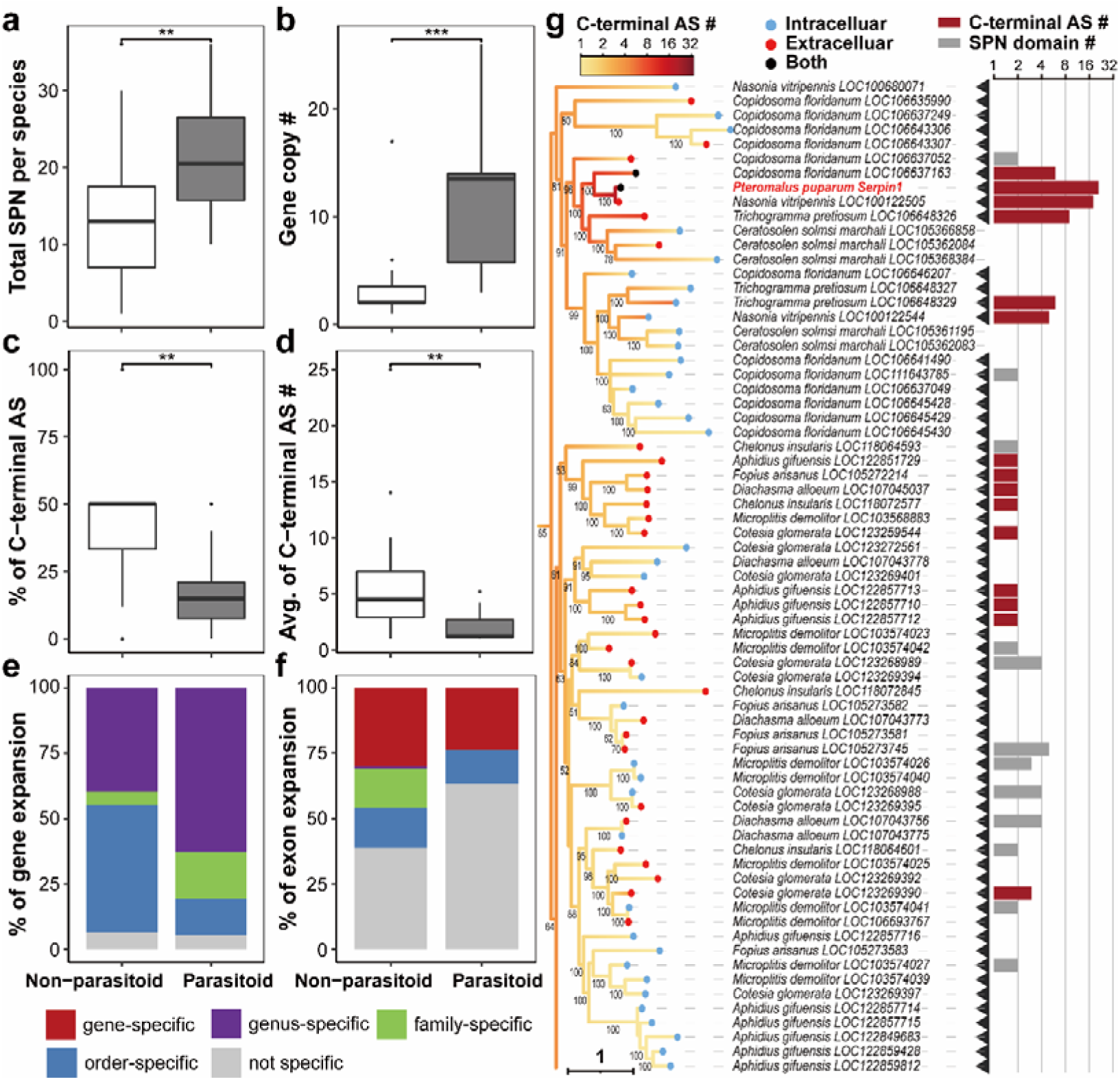
Parasitoid wasps employ fewer AS events in serpin, but PpSerpin-1 shows an expansion of the C-terminal AS number. (a-f) Comparison between parasitoid wasps and nonparasitoid hymenopterans of (a) total serpin protein per species, (b) gene copy number, (c) percentage of genes with C-terminal AS, (d) average C-terminal AS number, (e) distribution of gene expansion and (f) distribution of exon expansion. **(g)** Expansion of C-terminal AS number in PpSerpin-1. Black arrows indicate parasitoid wasps. The red and blue dots on branch terminals indicate the presence (extracellular) and absence (intracellular) of the signal peptide, respectively. Black dots on branch terminals indicate N-terminal AS with both extracellular and intracellular isoforms. The gradient color on branches indicates the estimated ancestral states of C-terminal AS numbers. The red bars indicate the log2 transformed numbers of C-terminal AS events. The gray bars indicate the log2 transformed numbers of serpin domains within a gene. *** *p* < 0.001, ** *p* < 0.01

However, PpSerpin-1 shows an expansion of AS number compared to other parasitoids. The C-terminal AS numbers of the PpSerpin-1 and *Nasonia vitripennis* LOC100122505 genes are increased compared to their homologous genes in the Chalcidoidea wasps, i.e., *Trichogramma pretiosum*, *Ceratosolen solmsi marchali*, and *Copidosoma floridanum* (Fig. 3g). For PpSerpin-1 alternative exons, H and X2 clustered together with the XP_008201831.1-specific exon of the *N. vitripennis* LOC100122505 gene as the closet outgroup, suggesting PpSerpin-1-specific exon expansion after divergence from the common ancestor with the *N. vitripennis* LOC100122505 gene, which is estimated to have occurred approximately 19 million years ago (MYA)^44^. Sixteen out of 21 alternative exons show clear orthologous relationships with alternative exons of the *N. vitripennis* LOC100122505 gene, suggesting that most alternative exons of PpSerpin-1 existed prior to the divergence between *Pteromalus* and *Nasonia*.

We then estimated the pairwise substitution rates of orthologous exons between the PpSerpin-1 and *N. vitripennis* LOC100122505 genes. Compared to constitutive exons, alternative exons show higher nonsynonymous substitution rates (dN) (Fig. S4b; MWUT; *W* = 15; *p* = 0.013) and lower synonymous substitution rates (dS) (Fig. S4c; MWUT; *W* = 84; *p* = 0.0061), resulting in higher dN/dS values (Fig. S4a; MWUT; *W* = 5; *p* = 5.1e^-4^). These results suggest accelerated protein evolution in alternative exons with lower substitution rates in synonymous sites, which can be important in the regulation of the AS process^1,2,45^.

### PpSerpin-1 is involved in the wasp’s immune response and recruited into both wasp venom and larval saliva

Next, we investigated the function of isoforms of PpSerpin-1. First, isoform-specific RT□CPCR confirmed the presence of all 21 C-terminal AS forms of PpSerpin-1 (Fig. 4a). These isoforms were more likely to be upregulated by the gram-positive bacterium *M. luteus* and the fungus *B. bassiana* than by PBS or the gram-negative bacterium *E. coli* (Fig. 4b; WSRT; all comparisons: *p* < 0.001), suggesting that PpSerpin-1 may be involved in the Toll pathway, which is preferentially activated by gram-positive bacteria and fungi^43^.

**Figure 4.**
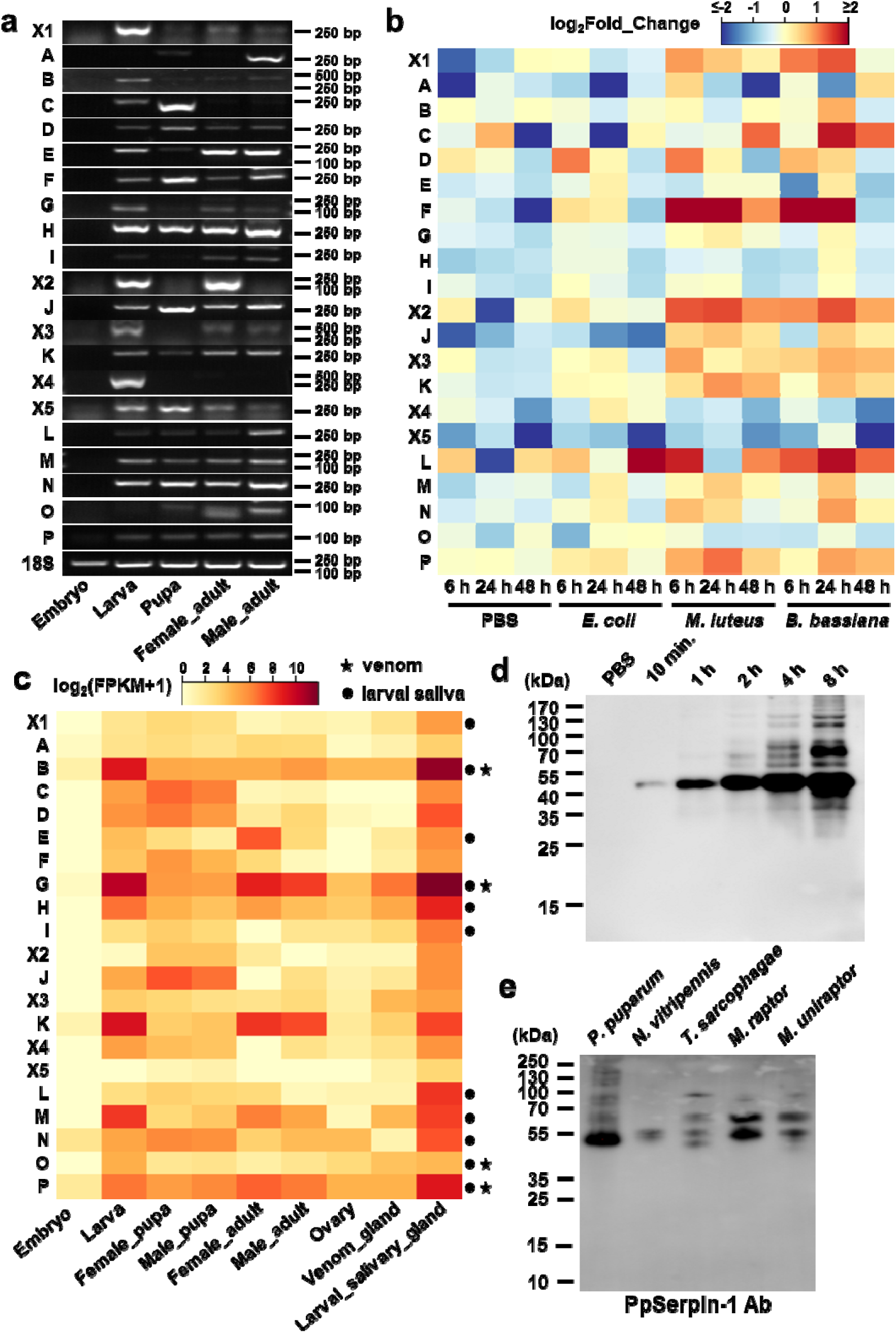
PpSerpin-1 isoforms are involved in the wasp’s immune response and parasitism. (a) Isoform-specific RT□CPCR of PpSerpin-1 isoforms. **(b)** Expression of PpSerpin-1 isoforms in response to injection of PBS, *E. coli*, *M. luteus* or *B. bassiana*. **(c)** Expression of PpSerpin-1 isoforms in different developmental stages and tissues. Black dots indicate proteomic identification in *P. puparum* larval saliva. Black stars indicate proteomic identification in venom of *P. puparum* female wasps. **(d)** Western blot detection of PpSerpin-1 proteins in PBS incubated with *P. puparum* larvae. **(e)** Western blot detection in larval saliva of *Nasonia vitripennis*, *Trichomalopsis sarcophagae*, *Muscidifurax raptor* and *M. uniraptor* using PpSerpin1 antibodies.

In addition, both RT□CPCR and transcriptomic results showed that some PpSerpin-1 isoforms were highly expressed in the larval stage (Fig. 4a, 4c), particularly in the larval salivary gland (Fig. 4c). We therefore hypothesize that some isoforms have been recruited into the larval saliva of *P. pupuarum*. Consistent with this hypothesis, PpSerpin-1 proteins were detected by Western blot using PpSerpin-1 antibodies in PBS incubated with *P. puparum* larva and in larval saliva of *P. puparum* (Fig. 4d). Protein homologs of PpSerpin-1 were also detected in the larval saliva from relatives of *P. puparum*, i.e., *N. vitripennis*, *Trichomalopsis sarcophagae*, *Muscidifurax raptor* and *M. uniraptor* (Fig. 4e). Moreover, we identified 11 isoforms of PpSerpin-1, i.e., X1, B, E, G, H, I, L, M, N, O and P, in the larval saliva of *P. puparum* by the protomic approach (Fig. 4c).

### PpSerpin-1 isoforms inhibit host melanization immunity

To investigate the function of PpSerpin-1, recombinant proteins of each isoform were produced (Fig. 5a). PpSerpin-1 isoforms A, G, O and P significantly inhibited hemolymph melanization of *P. rapae* (Fig. 5b; Dunnett’s test; all comparisons: *p* < 0.001) in a dose-dependent manner (Fig. 5c) and formed complexes with host hemolymph proteins in pull-down assays (Fig. 5d). Pull-down assays were also performed for the remaining isoforms, which were identified in wasp venom or larval saliva. PpSerpin-1B and H formed complexes with host hemolymph proteins, although they showed no inhibition of host melanazation (Fig. 5e).

**Figure 5.**
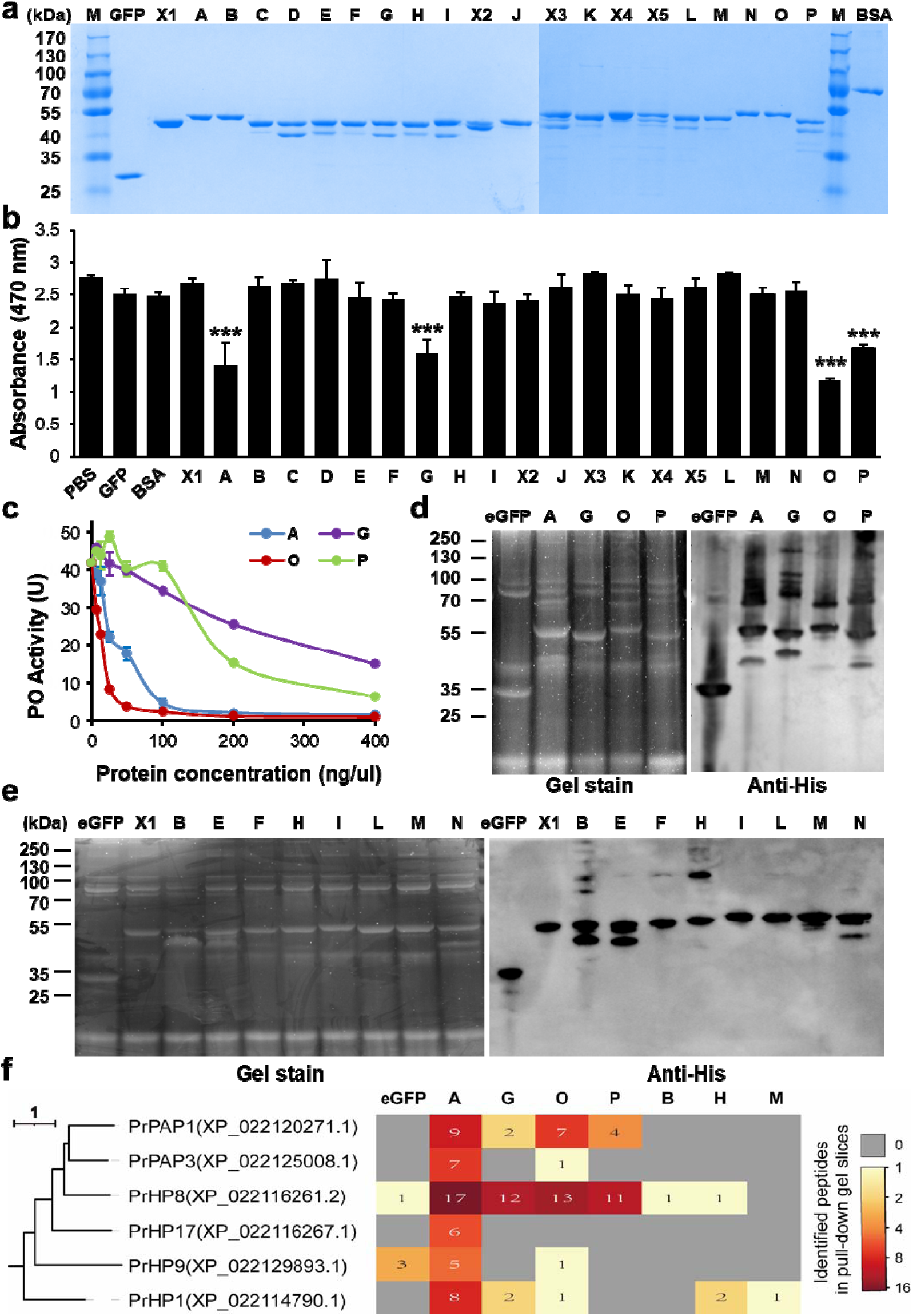
PpSerpin-1 isoforms inhibit host melanization and form complexes with host hemolymph proteins. (a) SDSCPAGE analyses of purified recombinant PpSerpin-1 isoform proteins. **(b)** Effect of PpSerpin-1 isoforms on PO activity of host *P. rapae* hemolyph. *** *p* < 0.001. **(c)** Dose-dependent suppression of host PO activity by PpSerpin-1A, G, O and P isoform proteins. **(d)** Pull-down assays of PpSerpin-1A, G, O and P isoform proteins from host hemolyph. **(e)** Pull-down assays of other PpSerpin-1 isoform proteins identified in venom or larval saliva of *P. puparum*. (f) Host hemolyph proteinases identified in pull-down samples of PpSerpin-1 isoforms. *Pieris rapae* hemolyph proteinases (PrHPs) are named based on their orthology with *Manduca sexta* hemolyph proteinases^46^. Pull-down samples of eGFP and PpSerpin-1M were used as negative controls.

To identify the targets of isoforms A, B, G, H, O, and P, we cut gel slices covering the complexes ranging from ∼60 to 100 kDa and identified 59, 9, 56, 29, 42, and 86 host proteins, respectively (Table S3). Among them, six identified proteins were serine proteases, i.e., PrPAP1, PrPAP3, PrHP8, PrHP17, PrHP and PrHP1 (Fig. 5f). The nomenclature of *Pieris rapae* hemolyph proteases (PrHPs) was based on their orthology with *Manduca sexta* hemolyph proteases^46^ (Fig. S5).

Among these protease candidates, PrPAP1 is *P. rapae* prophenoloxidase-activating proteinase 1 and critical for hemolymph melanization. The presence of PrPAP1 in pull-down samples of PpSerpin-1A, O and P was confirmed by Western blot using PrPAP1 antibodies (Fig. 6a). In vitro inhibitory assays showed that isoforms A, O and P but not G significantly inhibited the activity of recombinant PrPAP1 (Fig. 6b; Dunnett’s test, for A, O and P: *p* < 0.001; for G: *p* = 0.94). Consistent with this, isoforms A, O, and P but not G formed SDSCPAGE stable complexes with recombinant PrPAP1 (Fig. 6c, 6d and S6). Presumably, two identified peptides of PrPAP1 in the pull-down sample of PpSerpin-1G may be leaks from the neighboring pull-down samples of PpSerpin1-A and O. The stoichiometry of inhibition (SI) of PrPAP1 by PpSerpin1-A and P were 13.37 and 197.33, respectively (Fig. 6e, 6f), which were larger than the previously reported SI of 2.3 by PpSerpin-1O^22^. These results demonstrate that PpSerpin-1 isoforms A, O, and P suppress host melanization immunity by inhibiting PrPAP1.

**Figure 6.**
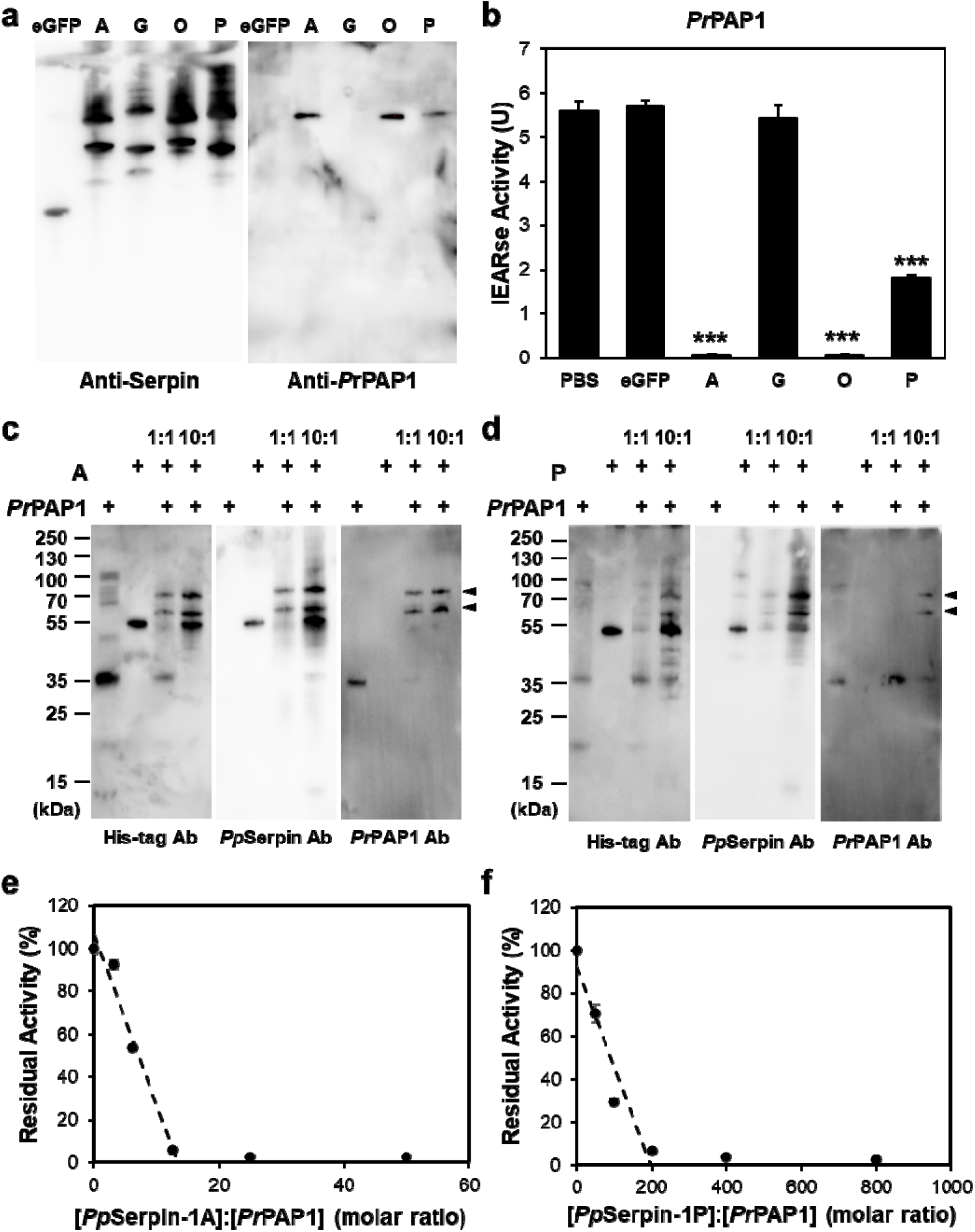
PpSerpin-1 isoforms A, O, and P but not G inhibit host PrPAP1. (a) Western blot detection of PrPAP1 in pull-down samples of PpSerpin-1A, G, O and P. **(b)** Effect of PpSerpin-1A, G, O and P on PrPAP1 activity. *** *p* < 0.001. **(c)** SDS-stable complex formation between PpSerpin-1A and PrPAP1. **(d)** SDS-stable complex formation between PpSerpin-1P and PrPAP1. **(e)** Stoichiometry of PrPAP1 inhibition by PpSerpin-1A. **(f)** Stoichiometry of PrPAP1 inhibition by PpSerpin-1P.

## Discussion

In this study, we report the divergent features of AS and GD in the evolution of insect serpins and their potential associations with the parasitic life strategy. We also find that PpSerpin-1 shows a number expansion of AS exons, which is involved in the wasp’s immune response and recruited to both wasp venom and larval saliva to suppress host immunity.

Using Illumina RNA-seq data, an exhaustive search was performed for AS isoforms of PpSerpin-1, but it is still possible that some isoforms may be missed. Long read sequencing, e.g., PacBio and Nanopore, may provide a more comprehensive and complete picture in future research^47^. In addition, proteomic identification also has limitations^48^. For example, PpSerpin-1A inhibits host melanization but was not identified in the venom or larval saliva of *P. puparum*. One possibility is that some isoform proteins were not detected by proteomics due to their low expression or short specific sequence regions. Conversely, several isoforms identified in wasp venom or larval saliva failed to form complexes with host hemolymph proteins. These PpSerpin-1 isoform proteins may target proteins out of the host hemolymph, e.g., in host hemocytes or gonad cells, inside host guts, or regulate proteases from wasp venom or larval saliva. Alternatively, the inconsistency between proteomic identification and functions of PpSerpin-1 isoforms may be simply due to leaky expression from AS regulation. More investigations are needed to test these hypotheses.

In addition to AS, GD and domain duplication jointly contribute to the protein diversity of insect serpins. Among them, AS and GD are the major sources of protein diversity. We observed an inverse relationship between the production of AS variants and gene family size across different species within a serpin clade. This is similar to previous reports that the production of AS variants was inversely correlated with the size of the gene family within the same species^9,10,49^. These results suggest negative regulatory mechanisms between AS and GD in the production of protein diversity.

AS is an efficient mechanism to generate significant protein diversity without requiring duplication and divergence of the entire gene^2,3,8^. In serpin AS genes, C-terminal AS tends to occur at the end of the hinge region with an extra nucleotide G. The serpin hinge region is critical to the conformational change and thus the inhibitory function of serpin^31,32^, while the extra nucleotide G may be important in the splicing process of AS. The splicing sites in AS serpins reflect the maximum reuse of sequence. Compared to non-AS genes, AS genes are more conserved in the constitutive region but more variable in the alternative region. In addition, in the comparison of PpSerpin-1 with its orthologous gene LOC100122505 in *N. vitripennis*, the alternative exons show significantly higher nonsynonymous substitution rates than the constitutive exons. Together, these results suggest that AS can allow rapid evolution of serpin proteins with relatively few sequence alternations while preserving most of the functional backbone.

On the other hand, the regulation of AS is often complicated^50^. For pairwise substitution rates between PpSerpin-1 and its orthologous gene LOC100122505 in *N. vitripennis*, alternative exons show significantly lower synonymous substitution rates than constitutive exons, suggesting that synonymous sites may be critical in the regulation of AS and thus under higher selective pressure in alternative exons^1,2,45^. In addition, the binding motifs of two transcription factors, dl and Dif, are enriched in the AS gene set compared to the non-AS gene set. Accumulated evidence suggests that transcription factors can be directly or indirectly involved in the regulation of AS, although the mechanisms are not fully understood^51,52^. As pre-mRNA splicing mainly takes place during transcription, extensive crosstalk has been reported between these two processes^51^. Binding motifs of dl and Dif are preferred on the intron region of the coding strand of AS genes. One possible explanation is that the transcription factors dl and Dif may regulate AS processes by binding intron regions of pre-mRNA.

To manipulate their hosts, parasitoid wasps often recruit effector genes, e.g., venom or larval salivary genes^14,15^. Serpin, a common component of parasitoid venom^17, 20–23, 33–41^, has also been reported in teratocytes of parasitoid wasps for host regulation^53,54^. Here, we further extend serpin recruitment to wasp larval saliva. A previous study also reported that *P. puparum* larval saliva inhibits host immunity^55^. In light of this, the higher numbers of serpin protein/domains in parasitoid wasps may be related to the recruitment of host effector genes. Another example of an AS venom gene is LOC122512947 from *Leptopilina heterotom* with 2 N-terminal and 20 C-terminal AS isoforms^34^. In addition, *Leptopilina boulardi* LbSPNy (ACQ83466.1) was reported as a venom non-AS gene to inhibit host melanization^41^. LbSPNy forms a *Leptopilina*-specific non-AS gene cluster with LOC122509667, 122505241, 12200434, 122502279, 122503460, and 122502269 of *L. heterotoma*, suggesting that LbSPNy likely originated from GD. Extensive GD of the serpin gene was also reported in the venom of *M. mediator*^21^. Our findings further show that the increased number of serpins in parasitoids results from more GD but less AS. This is consistent with the general observation that venom genes in parasitoid wasps and other venomous animals are more likely to originate from GD than AS^14,56^.

In PpSerpin-1, we observed a number expansion of alternative exons compared to other parasitoid serpin genes. Reconstructing the accurate evolutionary history of these exons is challenging, particularly for ancient expanded exons, due to their accelerated protein evolution and short sequences (44∼50 amino acids for PpSerpin-1 alternative regions). Phylogenetic tree analysis reveals that alternative exons of PpSerpin-1 H and X2 form a gene-specific cluster, suggesting a recent *Pteromalus*-specific exon duplication. However, the majority (16 out of 21) of PpSerpin-1 alternative exons show clear orthologous relationships with those of the *N. vitripennis* LOC100122505 gene. One possible explanation is that these exons underwent lineage-specific exon duplications before the divergence of *Pteromalus* and *Nasonia* (∼ 19 MYA^44^) and were retained in both species, probably serving conserved and essential functions. The expanded alternative exons in PpSerpin-1 may contribute to its ecological adaptation. Consistent with this hypothesis, we find that several isoforms of PpSerpin-1 are involved in the wasp’s immune response, have been recruited to both wasp venom and larval saliva, and suppress host immunity. However, no obvious expressional specialization, especially to venom glands, was observed for PpSerpin-1 isoforms. We speculate that accurate expressional regulation is an obstacle to the recruitment of host effector genes by AS, although AS may allow rapid evolution of protein sequences. In AS genes, differential expressional regulation between self and host manipulation functions may be difficult to evolve. On the other hand, expressional divergence of duplicated genes often occurs after location segregation of gene copies^57^, presumably due to changes in cis-regulatory regions. The positional linkage of alternative exons makes the expressional regulation of AS more challenging and requires more complicated regulatory mechanisms. In particular, high expressional specialization should be required to avoid self-harm for toxic virulent genes. This may explain why AS is less employed than GD in the recruitment of parasitoid host effector genes.

In summary, we show that both AS and GD contribute to the evolution of insect serpin with differential features. We report that a parasitoid serpin gene has evolved through exon number expansion of AS and show its involvement in wasp’s immunity and recruitment into the wasp’s venom and larval saliva to manipulate host immunity.

## Materials and Methods

### Insect rearing

Laboratory cultures of *P. puparum*, *N. vitripennis*, *T. sarcophagae*, *M. raptor* and *M. uniraptor* were maintained in *Drosophila* tubes at 25 °C with a photoperiod of 14:10 h (light:dark) as previously described^17,35^. Pupae of *Pieris rapae* were used as hosts for *P. puparum*, and pupae of *Sarchophaga bullata* were used as hosts for *N. vitripennis*, *T. sarcophagae*, *M. raptor* and *M. uniraptor*. Once emerged, *P. puparum* adult wasps were fed *ad libitum* with 20% (v/v) honey solution to lengthen their life span.

### Alternative isoform identification for the PpSerpin-1 gene

Using accumulated transcriptomic data^44^, sequencing reads were filtered by Trimmomatic v0.38^58^, mapped to the *P. puparum* genome by TopHat v2.0.12^59^, and assembled into transcripts using Cufflinks v2.2.1^59^. For exhaustive identification of PpSerpin-1 isoforms, we further manually identified isoforms using IGV browser v2.3.91 based on mapped reads^60^. The expression levels of PpSerpin-1 isoforms were estimated using Cufflink v2.2.1^59^.

### Protein structure prediction

After removing the single peptide, the protein structure of PpSerpin-1F was modeled using AlphaFold2^61,62^. Alignemnts for protein modeling were generated through MMSeqs2^63^ against the UniRef+Environmental database.

### Sequence fetch and feature analyses

Blastp^64^ was performed using the constitutive protein sequence of the PpSerpin1 gene against the Refseq_protein database on NCBI (accessed at Dec 2021). The organism was limited to “Insecta (taxid:50557)” with a maximum target sequence of 5000 and an e-value of 1e^-5^. One species per genus was selected as a representative to reduce sampling bias. GenBank files were retrieved for each gene using EFetch v0.2.2 (https://dataguide.nlm.nih.gov/edirect/efetch.html). Isoform sequences and splicing positions were extracted from GenBank files for each gene using homemade python scripts. For each gene, the number of N-terminal AS events was approximately estimated by counting the number of different sequences in the first 20 aa. Similarly, the number of C-terminal AS events was approximately estimated by counting the number of different sequences in the last 50 aa. Signal peptides were predicted using SignalP v5.0^65^. Serpin domains were annotated using the NCBI Batch Web CD-Search Tool (https://www.ncbi.nlm.nih.gov/Structure/bwrpsb/bwrpsb.cgi) against the “CDD 58235 PSSMs” database^66^ (accessed at Jan, 2022).

### Phylogenetic analyses

For gene phylogeny, the longest protein isoform was selected for each gene. Proteins were aligned using Mafft v7.310^67^. The gene tree was constructed using IQTree v2.2.0^68^ with an ultra bootstrap of 1000. The best-fit model (LG+R10) was automatically selected by built-in ModelFinder^69^ in IQTree. For exon phylogeny, annotated serpin domains were extracted with a C-terminal extending 30 aa or to the end of the sequences if less than 30 aa. For genes with C-terminal AS, only nonredundant C-terminal sequences were included. Serpin domain sequences were aligned using Mafft v7.310^67^. The tree was constructed using IQTree v2.2.0^68^. To reduce sampling bias, one species per genus was selected as a representative for visualization and subsequent statistical analyses. Ancestral character estimation was performed for the C-terminal AS number of each internal node using the fastAnc function in the R package “phytools” v1.2.0^70^. Trees were pruned by Newick Utilities v1.6^71^ and visualized on iTOL^72^.

### Expansion level analyses

For every pair of genes from the same species in the gene phylogeny, two genes are defined as species-specific expansion if all genes in the clade of the common ancestor of these two genes belong to the same species. If all genes in the clade of the common ancestor of these two genes from the same species belong to the same family, two genes are defined as family-specific expansion, and so on. Similarly, for every two alternative C-terminal exons from the same gene in the exon phylogeny, two exons are defined as gene-specific expansion if all exons in the clade of the common ancestor of these two exons belong to the same gene, and so on. Expansion levels were determined by homemade R scripts.

### WebLogo

Consensus sequence logos were created using WebLogo v3.7.9^73^. For protein alignments, conservation scores were estimated for each site using WebLogo v3.7.9^73^. Conservation scores of the corresponding positions in the reference sequence were extracted for comparisons. For the constitutive region of AS genes, the longest protein isoforms of each gene were selected as representatives. For the C-terminal alternative region of AS genes, all nonredundant sequences (based on the last 50 aa at the C-terminus) were used. Conservation scores were mapped to the PpSerpin-1F structure using PyMOL v2.5.0 (https://pymol.org).

### dN and dS estimation

For each orthologous exon pair between PpSerpin1 and LOC100122505 (the ortholog of PpSerpin1 in *N. vitripennis*), protein sequences were aligned using Mafft v7.310^67^ and then reverse translated to codon using PAL2NAL v14^74^. If a codon crossed the boundary of two exons, the entire codon was counted in the exon that contained more nucleotide bases of that codon. Pairwise dN and dS values were estimated using PAML v4.9j^75^.

### Motif scanning and enrichment analysis

Motif enrichment analyses were performed using SEA (Simple Enrichment Analysis) in MEME v5.5.0^76,77^. Gene sequences with C-terminal AS were set as primary sequences, and shuffled sequences or gene sequences without C-terminal AS were set as control sequences. *Drosophila melanogaster* “CIS-BP 2.00 single species DNA” or “CISBP-RNA single species RNA” was set as the motif database. Fisher’s exact tests were performed if the primary and control sequences had the same average length (within 0.01%); otherwise, binomial tests were performed^77^. Sequences of pomotor, exon and intron regions were extracted from GenBank files. Promoter regions were approximated by using the upstream sequences of the translation start sites. Exon regions were approximated using coding sequences by masking other regions using “N”. Motifs were detected using FIMO in MEME^76^. Shuffled sequences were created by fasta-shuffle-letters in MEME^76^.

### Specific RT**□**PCR

*Ptermoalus puparum* embryos (<10 hr after parasitism), larvae (combined 2–3 day larvae after parasitism), yellow pupae (mixed with female and male pupae), adult females and males (combined 1–5 day adults after emergence) were collected and rinsed with PBS. Total RNA was extracted using TRIzol reagent (Invitrogen, USA) according to the manufacturer’s protocol and then reverse transcribed using TransScript One-step gDNA Removal and cDNA Synthesis SuperMix (TransGen, Beijing, China) with random primers. Isoform-specific primers were designed to span constitutive exon 7 and alternative exon 8 using PerlPrimer v1.1.21^78^ and are listed in Table S1. PCRs were performed using TransTaq HiFi DNA Polymerase (TransGen, Beijing, China) with 20-35 cycles of amplification. PCR products were analyzed by electrophoresis on a 1% (g/mL) agarose gel and confirmed by Sanger sequencing (Sangon Biotech, Shanghai, China).

### Wasp larval saliva collection

Saliva of wasp larvae was collected as previously described with minor modifications^55^. Briefly, wasp larvae were collected by opening host pupae 3 days after parasitism. Larvae were rinsed with PBS and electrically stimulated using an acupuncture device (Hwato, China) with a frequency of 50 Hz and a current of 2.5 mA. Secreted saliva drops at larval mothparts were transferred into PBS using pipette tips. The protein concentration was determined using a Pierce BCA Protein Assay Kit (Thermo Scientific, USA).

### LC□MS/MS

Fifty micrograms of protein from wasp larval saliva was digested by trypsin using the filter-aided sample preparation (FASP) method. After desalting by a C18 cartridge, the digestion product was lyophilized and redissolved in 40 µl of 0.1% formic acid solution. LCCMS/MS was conducted on an Easy nLC HPLC system (Thermo Scientific, USA) with a flow rate of 300 nl/min followed by Q-Exactive (Thermo Finnigan, USA). The sample was loaded on a Thermo Scientific EASY column (5 µm, 2 cm × 100 µm, C18) and then separated on another Thermo Scientific EASY column (3 µm, 75 µm × 100 mm, C18). Buffer A was water with 0.1% formic acid, and buffer B was 84% acetonitrile with 0.1% formic acid. The columns were first equilibrated with 95% buffer A, then from 0% to 35% buffer B in 50 min, from 35% to 100% buffer B in 5 min, and finally 100% buffer B for 5 min. The charge-to-mass ratios of peptides and fractions of peptides were collected 20 times after every full scan. The resulting MS/MS spectra were searched against the *P. puparum* genome database using Mascot 2.2 in Proteome Discoverver. “Carbamidomethyl (C)” was set as a fixed modification. “Oxidation (M)” and “Acetyl (Protein N-term)” were set as variable modifications. The maximum number of missed cleavages was set as 2. FDR ≤ 0.01 was set to filter the protein identification. The same software and parameters were used for the reanalysis of *P. puparum* venom proteomic data^35^. This part was conducted by Shanghai Applied Protein Technology Co., Ltd. (Shanghai, China).

### Western blot

Protein was separated by electrophoresis on 12% SDSCPAGE and transferred to a PVDF (polyvinylidene difluoride) membrane at 100 mA for 2 h using a Mini-PROTEAN Tetra system (Bio-Rad, USA). For detection of PpSerpin1 isoform proteins or *Pr*PAP1, antibodies against *Pr*PAP1 and *Pp*Serpin1-O ^22^ (diluted 1:1000) were used as primary antibodies, followed by HRP (horseradish peroxidase)-conjugated goat anti-rabbit IgG antibody (Sigma Aldrich, Germany; diluted 1: 5000) as the secondary antibody. For detection of His-tagged proteins, THE™ His Tag mouse antibody (GenScript, Nanjin, China; diluted 1: 2000) was used as the primary antibody, followed by goat anti-mouse IgG antibody-HRP conjugate (Sigma Aldrich, Germany; diluted 1: 5000) as the secondary antibody. The membranes were detected using Pierce ECL Western Blotting Substrate ECL (Thermo Fisher, USA) and imaged by the Chemi Doc-It 600 Imaging System (UVP, Cambridge, UK).

### Recombinant protein expression and purification

For recombinant expression of PpSerpin1 isoforms, constitutive fragments without signal peptides and alternative fragments were separately amplified using TransTaq HiFi DNA Polymerase (TransGen, Beijing, China) and cloned into the pET-28a vector using the ClonExpress MultiS One Step Cloning Kit (Vazyme, Nanjing, China). Primers were designed using PerlPrimer V1.1.21 and are listed in Table S2. The linear vector pET-28a was generated by digestion with BamHI and XhoI (TaKaRa, Dalian, China). Recombinant plasmids were then transfected into *E. coli* BL21(DE3) Chemically Competent Cell (TransGen Biotech, Beijing, China) and confirmed by Sanger sequencing (Sangon Biotech, Shanghai, China). *E. coli* cells were grown in autoinduction medium^79^ containing 100 µg/µl kanamycin at 300 rpm and 20 °C for 48 h and then harvested by centrifugation at 12000 × g for 20 min. Recombinant protein was extracted using BugBuster Protein Extraction Reagent (Thermo Scientific, USA) and purified using cOmplete His-Tag Purification Resin (Roche, Switzerland) and His-Bind Purification kit (Novagen, USA) according to the manufacturer’s protocol. The concentration of purified protein was determined using a Pierce BCA Protein Assay Kit (Thermo Scientific, USA).

### Phenoloxidase activity assay

Plasma was harvested by scissors cutting the posterior gastropods of *P. rapae* larvae and diluted four times into ice-coled TBS buffer (20 mM Tris, 150 mM NaCl, pH=7.6). Cell-free hemolymph was obtained by centrifugation at 4 °C and 3000 × g for 10 min to remove hemocytes. To screen for the inhibitory activities of PpSerpin1 isoforms on host melanization, 5 µl of recombinant protein (0.2 µg/µl) was mixed with 10 µl of diluted *P. rapae* hemolymph in a 384-well plate. For each sample, 5 µl of elicitor (0.1 µg/µl *M. luteus*) and 5 µl of substrate solution (50 mM L-Dopa in PBS, pH=7.5) were first mixed and added to another 384-well plate, which was fixed upside down on the sample plate. By centrifuging these two oppositely fixed 384-well plates, the PPO (prophenoloxidase) cascade was activated in each well simultaneously. Plates were measured at A470 and 25 °C every 5 min for 2 h using a Varioskan Flash multimode reader (Thermo Scientific, USA). For dose-dependent assays, 5 µl of recombinant protein was mixed with 15 µl of diluted *P. rapae* hemolymph and 5 µl of elicitor (0.1 µg/µl *M. luteus*). After incubation at 25 °C for ∼10 min, 800 µl of substrate solution (20 mM Dopa in PBS, pH = 6.5) was added. Samples (200 µl) were monitored at A470 in 96-well plates for 20 min using a Varioskan Flash multimode reader (Thermo Scientific, USA). One unit of PO activity was defined as 0.001 △ A470/min.

### *Pr*PAP1 amidase activity assay

Recombinant *Pr*PAP1Xa was secretively expressed using the Bac-to-Bac™ Baculovirus Expression System (Invitrogen, USA) as previously described^22^. Unexpectedly, *Pr*PAP1Xa was activated for unknown reasons. For inhibitory assays, recombinant proteins of PpSerpin1 isoforms were incubated with activated *Pr*PAP1 at room temperature for 10 min. After adding 200 μl of 50 μM acetyl-Ile-Glu-Ala-Arg-p-nitroanilide (IEARpNA) in TBS (100 mM Tris, 100 mM NaCl, 5 mM CaCl_2_, pH = 8.0), residual amidase activities were measured at A405 for 20 min using a Varioskan Flash multimode reader (Thermo Scientific, USA). One unit of amidase activity was defined as 0.00001 △A405/min.

### Pull-down assay

Recombinant protein of PpSerpin1 isoforms (10 μg) was mixed with 1 ml diluted *P. rapae* hemolymph, 50 μl saturated PTU and 100 μl *M. luteus* (1 μg/μl) and incubated on a rotator overnight at 4 °C. After centrifugation at 12000 g and 4 °C for 20 min, the supernatant was further incubated with 25 μl of cOmplete His-tag purification resins (Roche, Switzerland) at 4 °C for 2 h. After washing three times with 300 μl washing buffer (1 M NaCl, 120 mM imidazole, 40 mM Tris-HCl, pH 7.9), proteins were eluted with 50 μl of elution buffer (1 M imidazole, 0.5 M NaCl, 20 mM Tris-HCl, pH 7.9) and subjected to SDSCPAGE followed by Lumitein Protein Gel Stain (Biotium, Hayward, CA, USA) and immunoblotting. To identify potential targets in pull-down complexes, gels were cut between 60 and 100 kDa. Gel slices were then in-gel digested by trypsin at 37 °C for 20 h. After desalting and lyophilization, the enzymatic product was redissolved in 0.1% formic acid solution and subjected to LCCMS/MS. The parameters used were the same as for the above, except specifically mentioned. The gradient was 1 h. Annotated proteins from *Pieris rapae* genome assembly GCF_001856805.1 were set as the search database. “Oxidation (M)” was set as a variable modification. Protein identification was filtered by proteins with at least two peptides identified. Gel digestion and proteomic identification were conducted by Shanghai Applied Protein Technology Co., Ltd. (Shanghai, China).

### Statistics

All statistical analyses were performed in R v4.1.2. Differences between Spearman correlations were tested using the Hittner2003 method in the R package “cocor” v1.1.4^80^. For independent contrasts, the phylogenetic species tree was generated by phyloT v2 (https://phylot.biobyte.de/). The branch length was estimated by compute.brlen in the R package “ape” v5.6.2^81^ using Grafen’s (1989) methods^82^. Independent contrasts were conducted using the “pic” function in the R package “ape”^81^. Correlations through oringins were estimated for independent contrasts using the R package “picante” v1.8.2^83^.

### Data availability

Alignments, trees and iTOL annotation files are publicly available on FigShare (https://doi.org/10.6084/m9.figshare.21545598.v1).

## Acknowledgments

This work was supported by the Key Program of National Natural Science Foundation of China (NSFC) (grant no. 31830074 to G.Y.Y.), NSFC (grant no. 31701843 to Z.C.Y.), the Regional Joint Fund for Innovation and Development of NSFC (Grant no. U21A20225 to G.Y.Y.), NSFC (grant no. 32072480 to Q.F. and no. 3200170366 to L.Y.), Hainan Province Natural Science Foundation (Grant no. 323QN262 to Y.Z.C.), China Postdoctoral Science Foundation (Grant no. 2021M700125 to X.H.Y.), Zhejiang Provincial Natural Science Foundation of China (Grant No. LTGN23C140001 to F. Q.), Major International (Regional) Joint Research Project of NSFC (Grant no. 31620103915 to G.Y.Y.). We thank Dr. John H. Werren (University of Rochester, USA) for providing *N. vitripennis*, *T. sarcophagae*, *M. raptor* and *M. uniraptor* species, as well as supervising Z.C.Y. in sampling their saliva. We also thank John H. Werren for his comments and suggestions on the manuscript.

## Author Contributions

G.Y.Y., F.Q., and Z.C.Y. conceptualized and designed the research; Z.C.Y. and F.Q. carried out the bioinformatics analysis; Z.C.Y. conducted the majority of the experiments; L.Y. performed immune induction and proteomics of larval saliva in *Pteromalus puparum*; S.X. performed recombinant expression of PpSerpin-1 proteins; J.L.W. performed RT□CPCR; Y.Z.C. and F.Q. interpreted the results; Z.C.Y. and G.Y.Y. wrote and revised the manuscript.

