## Supplemental figures for "A parasitoid serpin gene that disrupts host immunity shows adaptive evolution of alternative splicing"

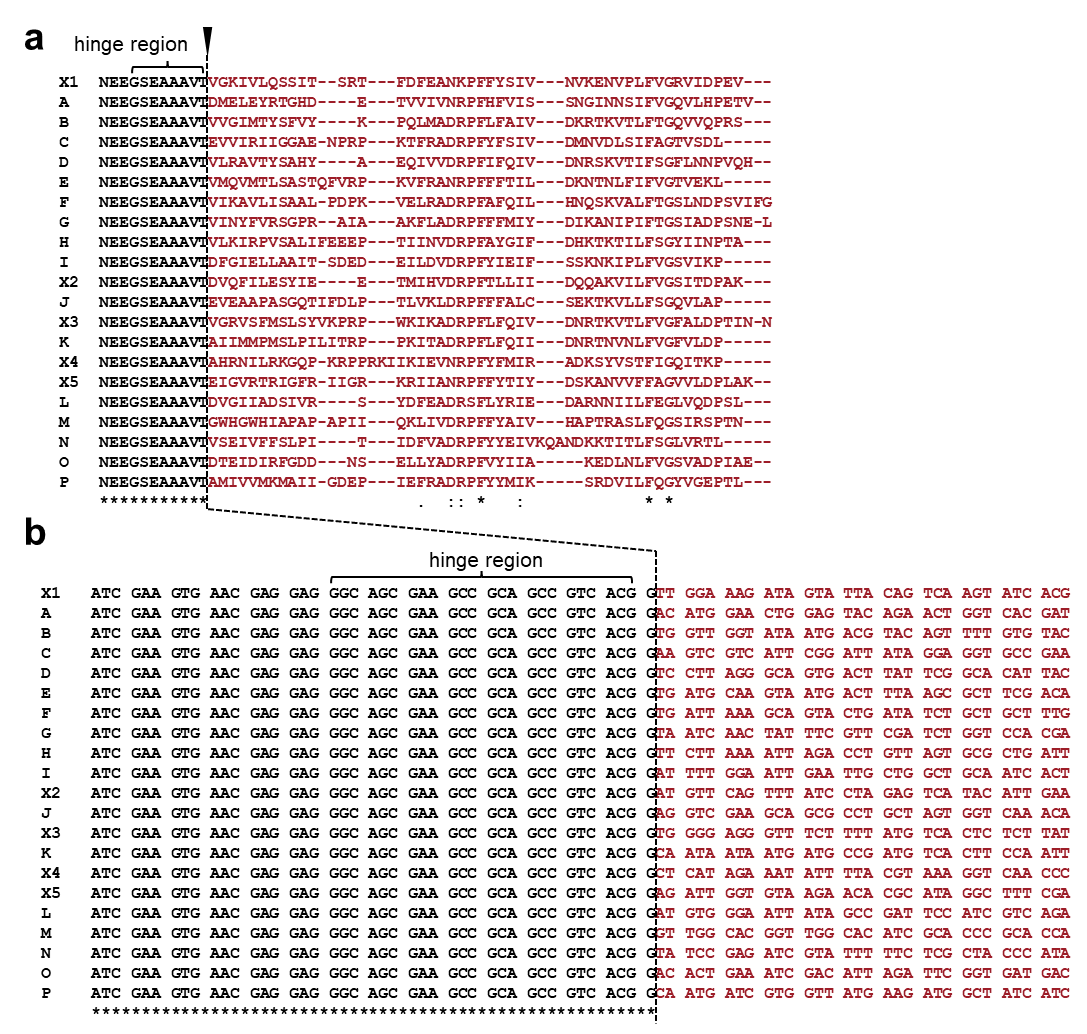

**Figure S1. Protein and codon alignments at the C-terminal alternative splicing site in the PpSerpin1 gene.** The black arrow indicates the splicing site. Curly brackets indicate the hinge region.

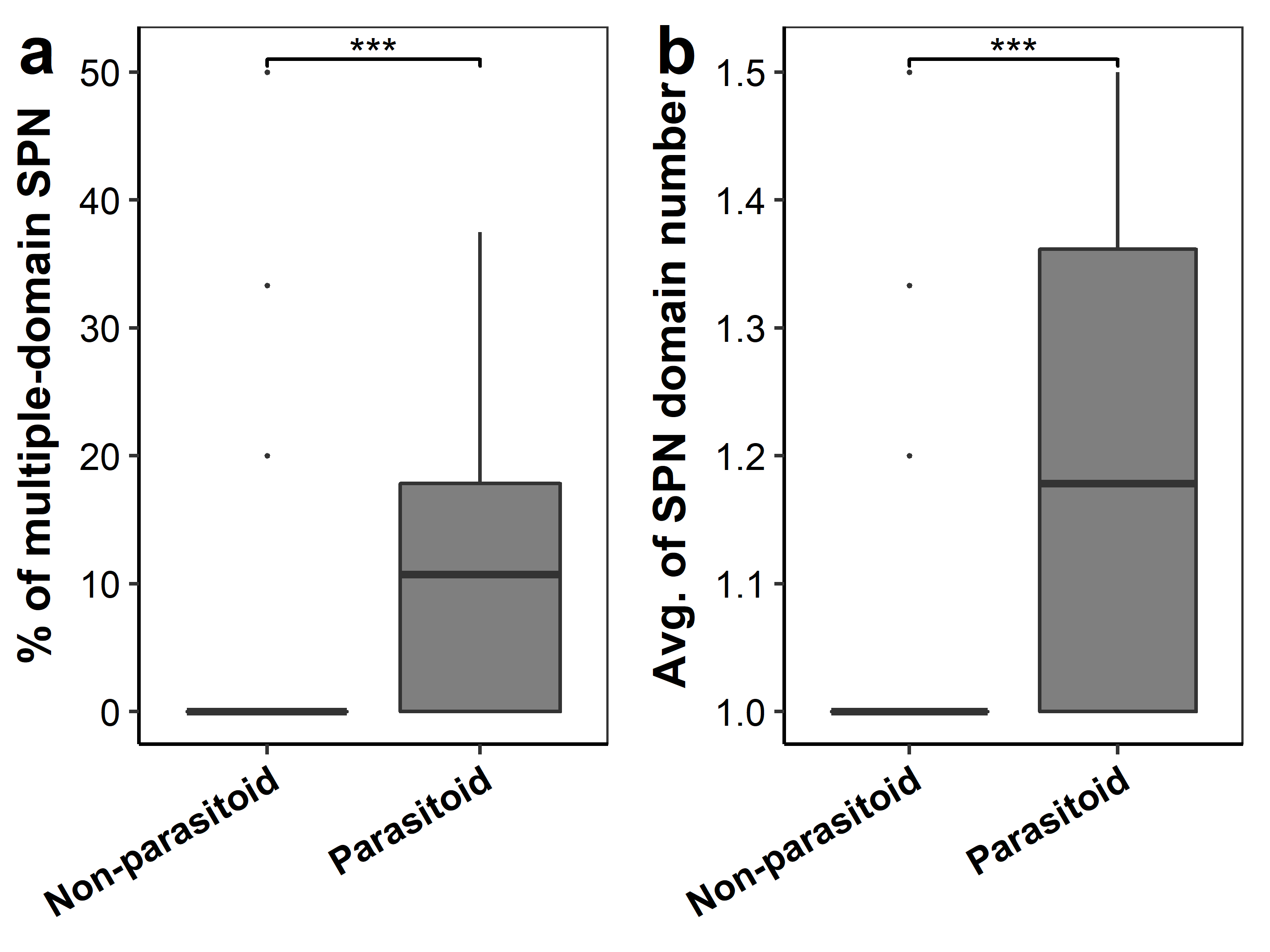

**Figure S2. Comparison of multiple-domain serpin between parasitoid wasps and non-parasitoid hymenopterans.** SPN, serpin.

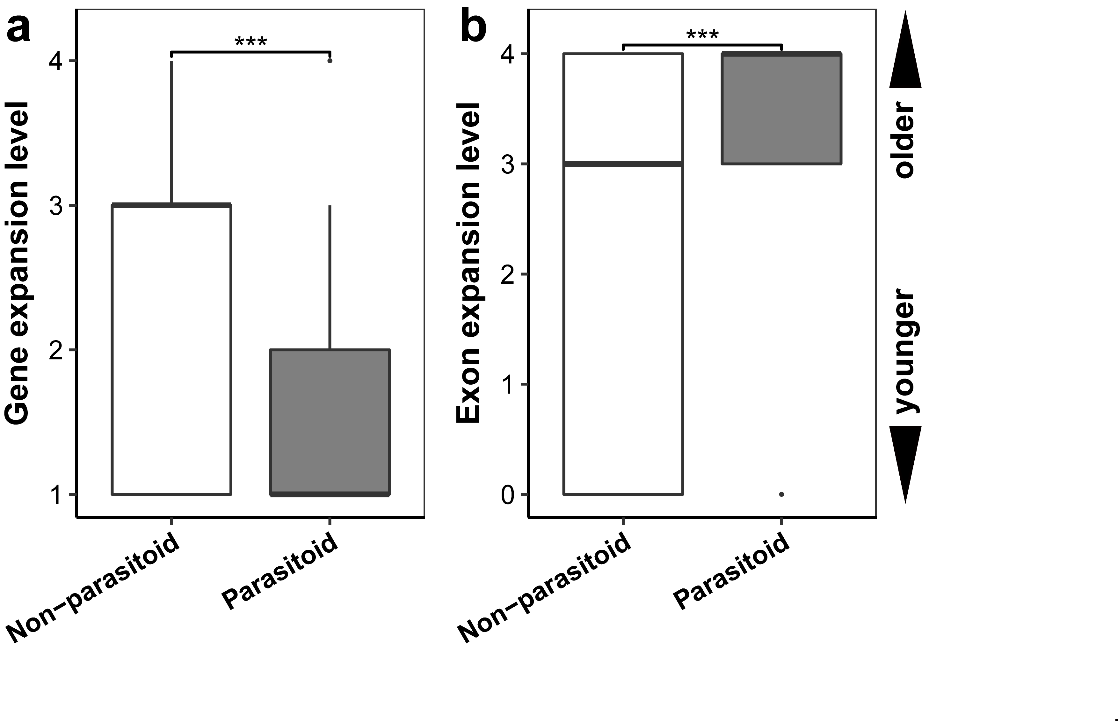

**Figure S3. Comparison of gene and exon expansion between parasitoid wasps and non-parasitoid hymenopterans.** Expansion levels were compared using a ranking method. 0 indicates gene-specific expansion; 1 indicates genus-specific expansion; 2 indicates family-specific expansion; 3 indicates order-specific expansion; 4 indicates not specific. Smaller rank numbers mean younger expansions.

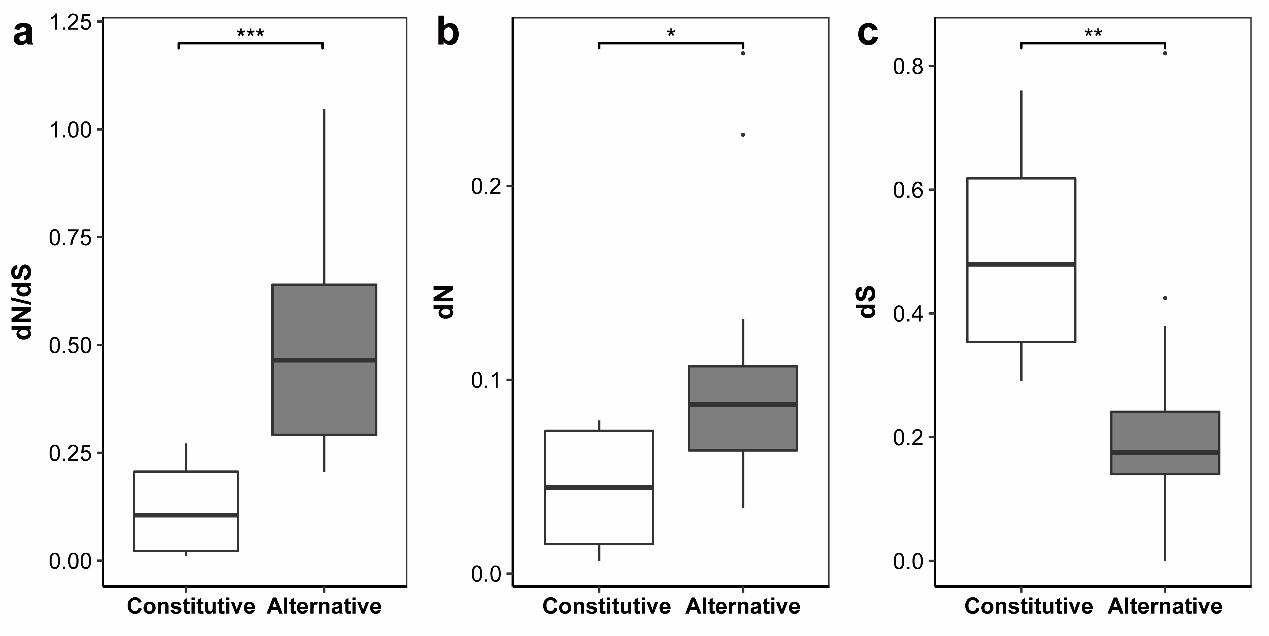

**Figure S4. Comparison of dN/dS between shared and alternative exons.** Constitutive exons include exons 2-7, and alternative exons include 8B, 8C, 8D, 8E, 8F, 8G, 8H, 8I, 8J, 8K, 8X4, 8X5, 8L, 8N, 8O, and 8P.

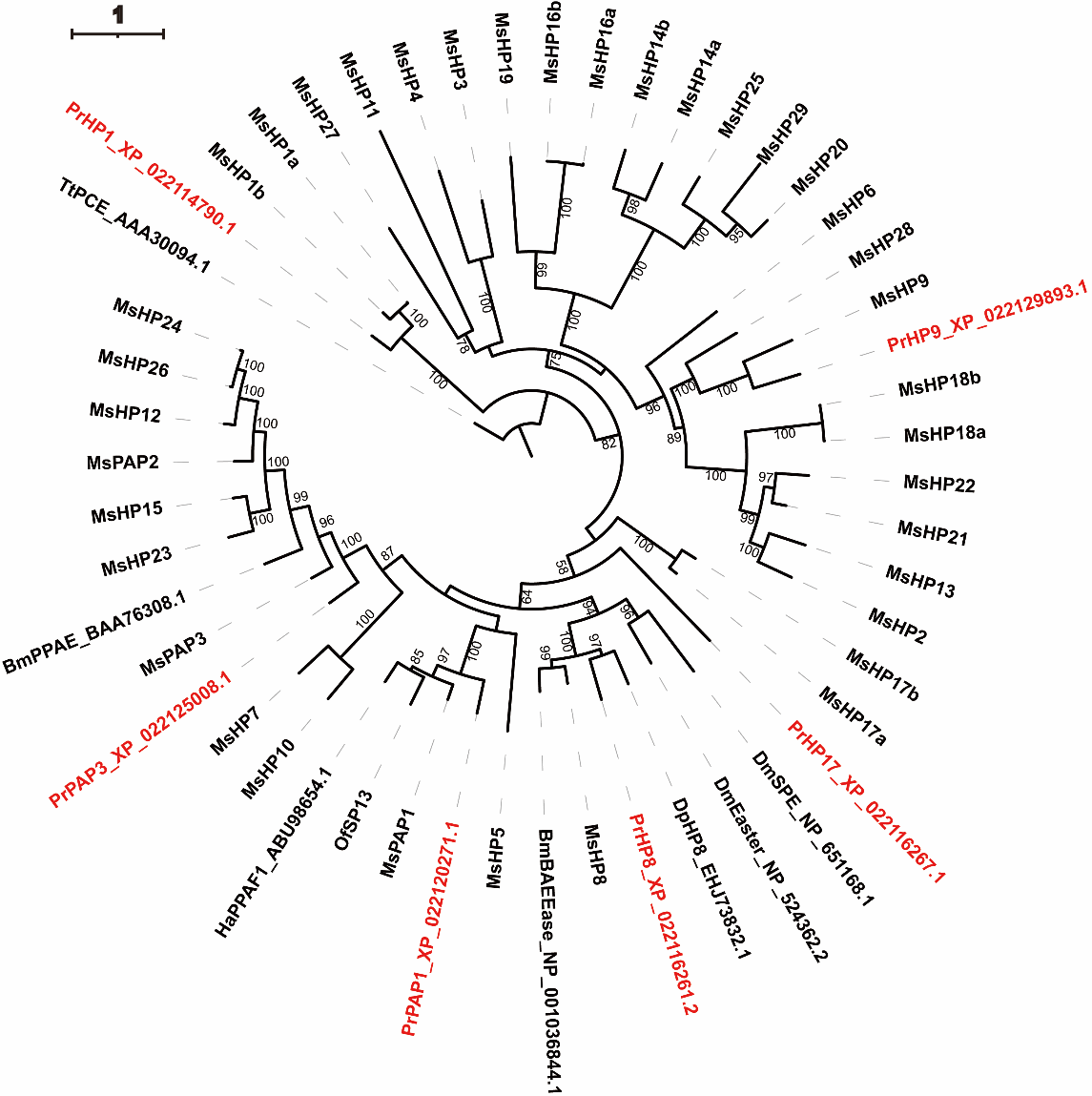

**Figure S5. Phylogenetic tree of PrHPs in pull-down samples of PpSerpin-1 isoform proteins.** Sequences of MsHP and MsPAP were retrieved from Cao et al. 2015 paper. Sequence, alignment and tree files can be downloaded at FigShare (https://doi.org/10.6084/m9.figshare.21545598.v1).

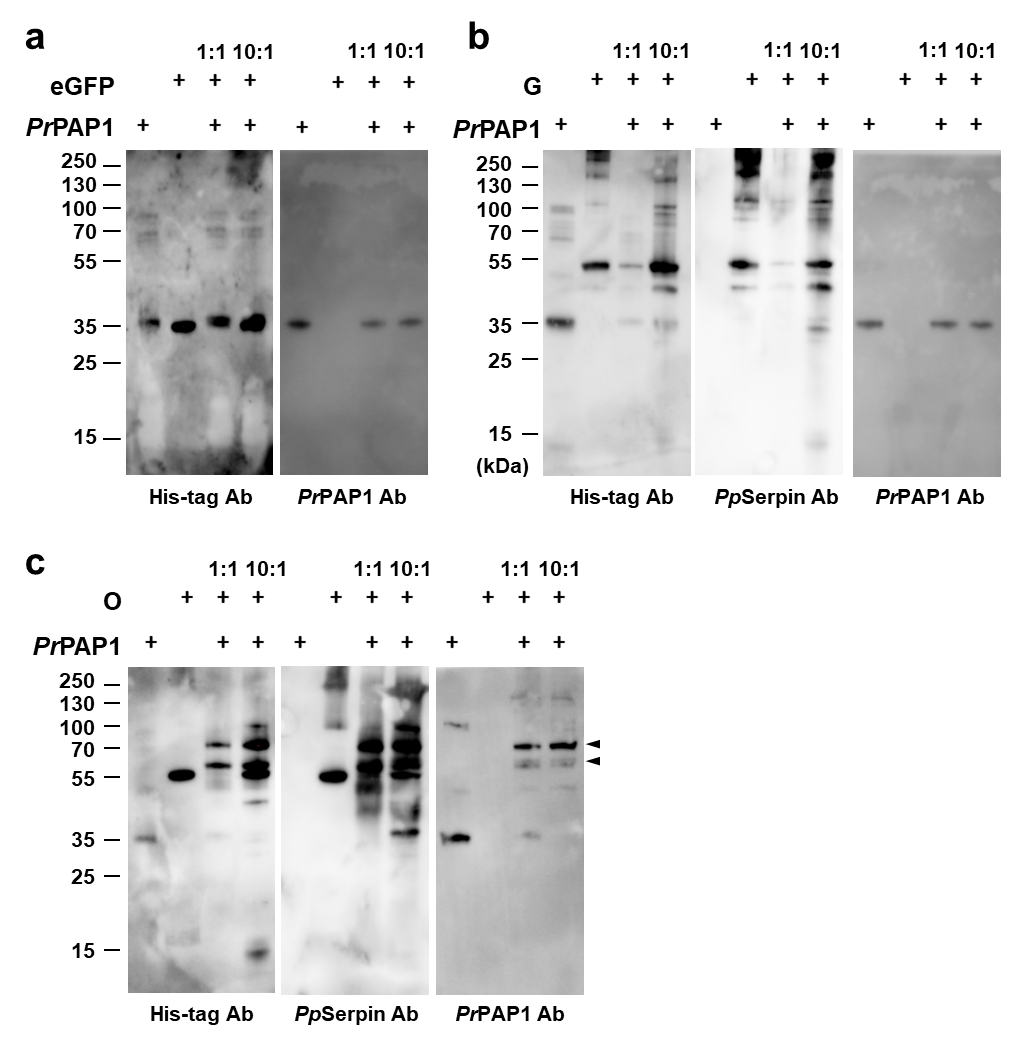

**Figure S6. Interaction of *Pr*PAP1 with eGFP, G and O.** Black arrows indicate complexes formed for PpSerpin1 isoforms with PrPAP1.
